# TNF-α induces reactivation of human cytomegalovirus independently of myeloid cell differentiation following post-transcriptional establishment of latency

**DOI:** 10.1101/298547

**Authors:** Eleonora Forte, Suchitra Swaminathan, Mark W. Schroeder, Jeong Yeon Kim, Scott S. Terhune, Mary Hummel

## Abstract

We used the Kasumi-3 model to study HCMV latency and reactivation in myeloid progenitor cells. Kasumi-3 cells were infected with HCMV strain TB40/E*wt*-GFP, flow sorted for GFP+ cells, and cultured for various times to monitor establishment of latency, as judged by repression of viral gene expression (RNA/DNA ratio) and loss of virus production. We found that, in the vast majority of cells, latency was established post-transcriptionally in the GFP+ infected cells: transcription was initially turned on, and then turned off. We also found that some of the GFP-cells were infected, suggesting that latency might be established in these cells at the outset of infection. We were not able to test this hypothesis because some GFP-cells expressed lytic genes, and thus, it was not possible to separate them from GFP-quiescent cells. In addition, we found that the pattern of expression of lytic genes that have been associated with latency, including UL138, US28, and RNA2.7, was the same as that of other lytic genes, indicating that there was no preferential expression of these genes once latency is established. We confirmed previous studies showing that TNF-α induced reactivation of infectious virus, and by analyzing expression of the progenitor cell marker CD34 as well as myeloid cell differentiation markers in IE+ cells after treatment with TNF-α, we showed that TNF-α induced transcriptional reactivation of IE gene expression independently of differentiation. TNF-α-mediated reactivation in Kasumi-3 cells was correlated with activation of NF-κB, KAP-1 and ATM.

**IMPORTANCE:** HCMV is an important human pathogen that establishes lifelong latent infection in myeloid progenitor cells, and reactivates frequently to cause significant disease in immunocompromised people. Our observation that viral gene expression is first turned on, and then turned off to establish latency suggests that there is a host defense, which may be myeloid-specific, responsible for transcriptional silencing of viral gene expression. Our observation that TNF-α induces reactivation independently of differentiation provides insight into molecular mechanisms that control reactivation.

## INTRODUCTION

HCMV is a ubiquitous human pathogen of the beta herpesvirus family. Infection is transmitted through contact with body fluids, including saliva, urine, blood, and genital secretions. Hematogenous spread of the virus leads to a systemic infection, with multiple cell types infected in many organs. In immunocompetent hosts, primary infection is typically sub-clinical, and is resolved by activation of innate and adaptive immunity. However, latently infected cells, which carry viral DNA, but do not produce infectious virus, are able to escape immune surveillance, and can persist for the life of the host.

Under the appropriate conditions, the virus in these cells wakes up from its dormant state to reactivate the infectious cycle. In immunocompetent hosts, viral replication is controlled by the immune response, but reactivation of latent virus can be a significant infectious complication in immunocompromised hosts, such as recipients of solid organ or bone marrow transplants. Reactivation of CMV in solid organ transplant recipients is associated with increased risk of CMV disease, acute and chronic allograft rejection, infection with other opportunistic pathogens, graft failure, and death (1). Risk factors for reactivation include CMV serostatus of the donor (D) and recipient (R), with the highest risk in the D+/R-combination, specific immunosuppression protocols, and inflammatory conditions associated with high TNF-α secretion, including allograft rejection and sepsis (1).

The cell type harboring latent CMV DNA in solid organs has not been clearly identified. However, it is known that latent virus is transmitted through blood transfusion, and monocytes in the blood and hematopoietic progenitor cells (HPCs) in the bone marrow are sites of latency in vivo (2-10). Experimental models have shown that these cells are less permissive to lytic replication, and that they support a latent infection (11-16). Cell type-specific establishment of latency is thought to be due to a combination of host and viral factors. Infection activates a host intrinsic immune response, which recognizes viral DNA invading the nucleus and silences viral gene expression at the outset of infection through heterochromatinization of viral genomes (13, 17-26). Factors present in the viral particle, including the tegument protein pp71, enter the cell upon infection, and counteract this host defense response to activate viral gene expression. In cells that support latency, pp71 is sequestered in the cytoplasm and is therefore unable to perform this function (26-28). Differentiation of myeloid cells to dendritic cells increases permissiveness of these cells to infection and also induces reactivation of both naturally and experimentally latently infected cells (12, 13, 15, 20, 29-32). Thus, the generally accepted paradigm is that myeloid progenitor cells are not permissive to infection due to heterochromatinization and transcriptional repression of invading viral genomes. The process of myeloid cell differentiation is thought to change the balance of repressive and activating cellular factors that control both viral and cellular transcription, resulting in changes to the viral epigenome that activate viral gene expression (17, 33, 34).

However, some recent observations suggest that the current paradigm may be in need of revision. First, several investigators have demonstrated that, although myeloid progenitors may be less permissive than other cell types, some expression of viral lytic genes is detectable at early times post-infection in various models of latency, including primary CD34+ HPCs, Kasumi-3 cells, a CD34+ myeloblastic cell line derived from a patient with acute myeloid leukemia (AML) (35), which is a tractable model for CMV latency and reactivation (27, 36, 37), and primary CD14+ peripheral blood mononuclear cells (11, 12, 36-43). Expression of these genes is lost as latency is established. These observations suggest that latency is not necessarily established at the outset of infection, but may also be established following activation of viral transcription.

Second, differentiation of myeloid cells into dendritic cells *in vivo* occurs in the context of infection and inflammation (44), and all of the physiologically relevant differentiation factors that have been used to induce reactivation of CMV in primary hematopoietic cells, including GM-CSF (45), TNF-α, IL-6, and LPS, are mediators of inflammation. Thus, it is difficult to distinguish the roles of inflammation versus differentiation in these models. In contrast, Kasumi-3 cells, like other transformed cell lines, are refractory to normal physiological differentiation cues, and therefore can be used to dissect the contributions of these two pathways. Here, we have used the Kasumi-3 model to further investigate these issues.

## RESULTS

### Latency is established after activation of transcription

The time at which latency is established following infection with HCMV has not been clearly determined. Some previous studies have shown that primary CD34+ myeloid progenitor cells do not express lytic viral genes upon infection, but rather, that viral genomes in these cells are inactivated upon entry and remain quiescent until they receive signals that induce them to differentiate (26, 46). However, other studies have shown that both primary CD34+ cells and Kasumi-3 cells express an SV40 promoter-driven GFP reporter, as well as viral genes at early times post-infection (11, 36-41), and that latency can be established in the GFP+ population following cell sorting (11, 36, 38, 40, 41, 47). To investigate the question of timing further, we infected Kasumi-3 cells with the TB40/E*wt*-GFP strain of HCMV (48), and analyzed the cells by FACS after 24 hr (Fig. 1A). GFP expression was observed in 20-50% percent of cells (Fig. 1B), as previously reported (36). Following inactivation of input virus, we purified the GFP+ population by cell sorting and analyzed viral RNA expression, DNA copy number, and production of infectious virus over the course of infection. Latency has been operationally defined as the lack of virus production in cells that carry viral DNA with the capacity to reactivate (49). Production of infectious virus was detectable at 4 dpi and peaked at 8 dpi, but was no longer detectable at 11dpi (Fig. 1C).

**Fig. 1.**
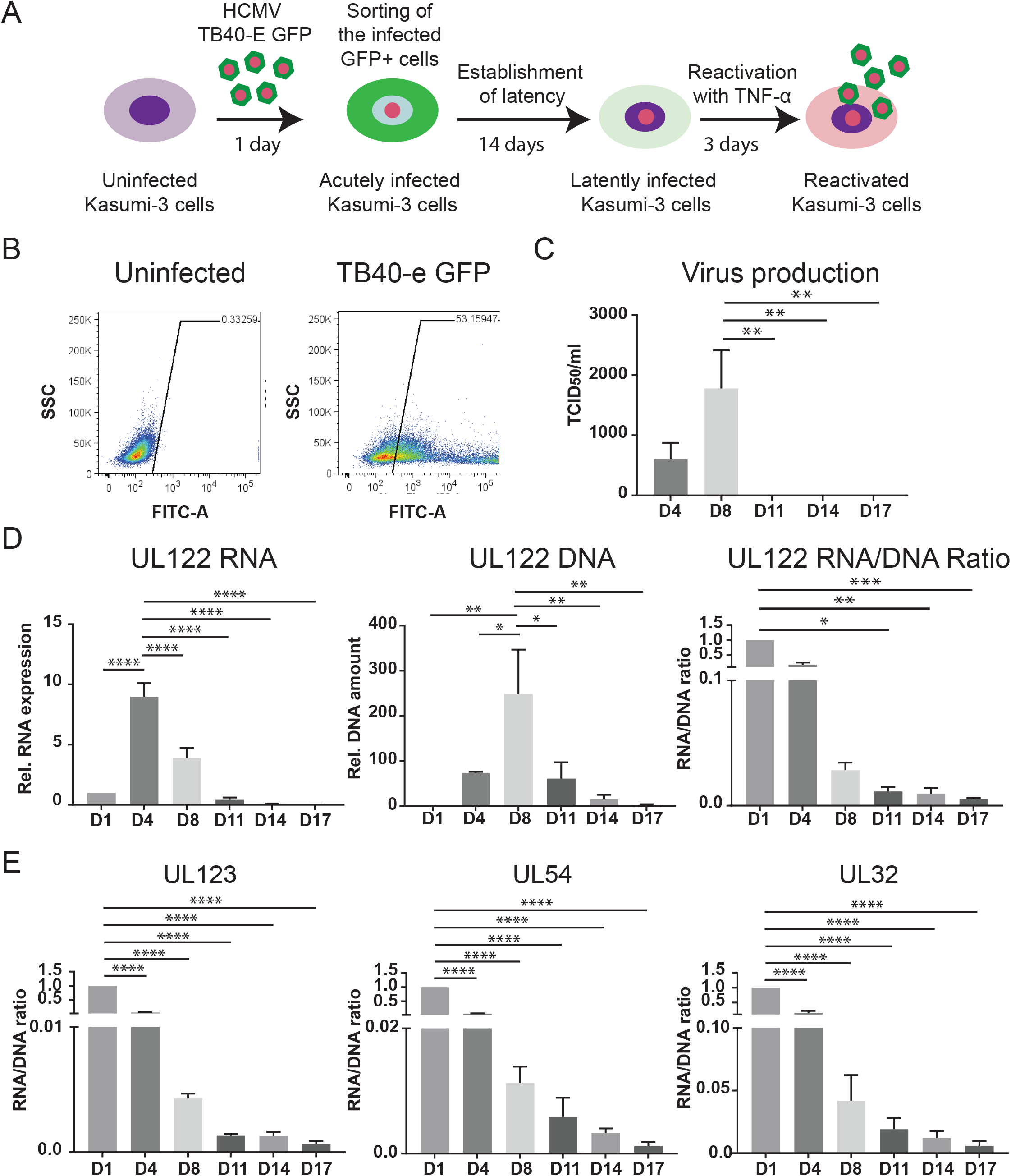
HCMV latency is established after activation of transcription at 14 days post infection. (A) Schematic outlining the infection model used for studies of latency and reactivation in Kasumi-3 cells. GFP+ infected cells were purified by flow cytometry at 1 dpi. On day 14, latently infected cells were treated with TNF-α for 3 days to induce reactivation. (B) Representative FACS analysis of GFP expression in Kasumi-3 infected cells at 1 dpi compared to uninfected cells. (C) Release of viral particles into the media was measured by a TCID_50_ assay on MRC-5 cells after 2 weeks. (D) UL122 mRNA expression and DNA amount were analyzed by at the indicated times post infection and expressed relative to D1 after normalization to GAPDH or RNAseP. (E) RNA/DNA ratios of UL123, UL54, and UL32 over the course of infection. For Panels C to E, statistical significance was calculated by a one-way ANOVA with a Dunnett’s multiple comparison test (n=4). The error bars represent standard error of the mean (SEM) and the asterisks indicate p-values (* P≤ 0.05; ** P≤ 0.01; *** P≤ 0.001; **** P ≤ 0.0001) calculated by the comparison to the peak (D4 for RNA, D8 for DNA and virus, and D1 for the RNA/DNA ratio).

A second characteristic of latency is transcriptional repression of genes involved in lytic replication. We therefore analyzed expression of viral genes representative of the immediate early (IE-2, UL122 and IE-1, UL123), early (viral DNA polymerase UL54), and late (UL32) phases of the viral life cycle. Expression of these genes peaked at 4 dpi and then fell throughout the course of infection. Viral DNA copy number also increased following infection, peaked 4-8 dpi, and then decreased significantly (Fig. 1D and Fig. S1). Because both viral DNA and RNA copy number dropped over the course of infection, analysis of RNA expression alone does not indicate repression of viral transcription. We therefore calculated the ratio of viral RNA to DNA. Relative to 1 dpi, this ratio fell for all genes from 4 to 14 dpi (Fig. 1D and E). TNF-α induced reactivation of infectious virus in these cells (see below). It is important to note that, because the cells were sorted for GFP expression to obtain a pure population of infected cells in which viral genomes were transcriptionally active, these results show that latency can be established by shutting off gene expression after it has been turned on. We have introduced the term post-transcriptional latency to describe this state.

### Some GFP-cells contain viral DNA and express lytic transcripts

Although our studies indicate that latency can be established post-transcriptionally, we wondered whether there might be a second population of cells in which latency is established at the outset of infection. These cells would carry viral DNA, but would not express GFP or viral genes. To test this hypothesis, we analyzed both the GFP-positive and GFP-negative populations for the presence of viral DNA at 1 dpi. To preclude cross-contamination of these fractions, the gates for sorting were set far apart. qPCR analysis revealed that, although the vast majority of the DNA was present in the GFP+ cells, some DNA was detectable in the GFP-population (Fig. 2C). We then analyzed the RNA from these cells for expression of immediate early, early, and late genes (Fig. 2D). Expression of immediate early (IE-2, UL122) RNA was detectable in the GFP-population, albeit at a lower level than the GFP+ cells. When we calculated the ratio of UL122 RNA to DNA, we found that there was no difference between the GFP+ and GFP-cells, indicating that transcription of the IE genes was equally active in these two populations at this time (Fig. 2E). However, when we analyzed the RNA/DNA ratios of early (UL54) and late (UL32) genes, we found that this ratio was significantly lower in the GFP-cells (Fig. 2E), indicating that these classes of genes were transcriptionally repressed in the GFP-cells. We then analyzed the GFP-cells for changes in GFP expression, viral RNA expression and DNA copy number over time.

**Fig. 2.**
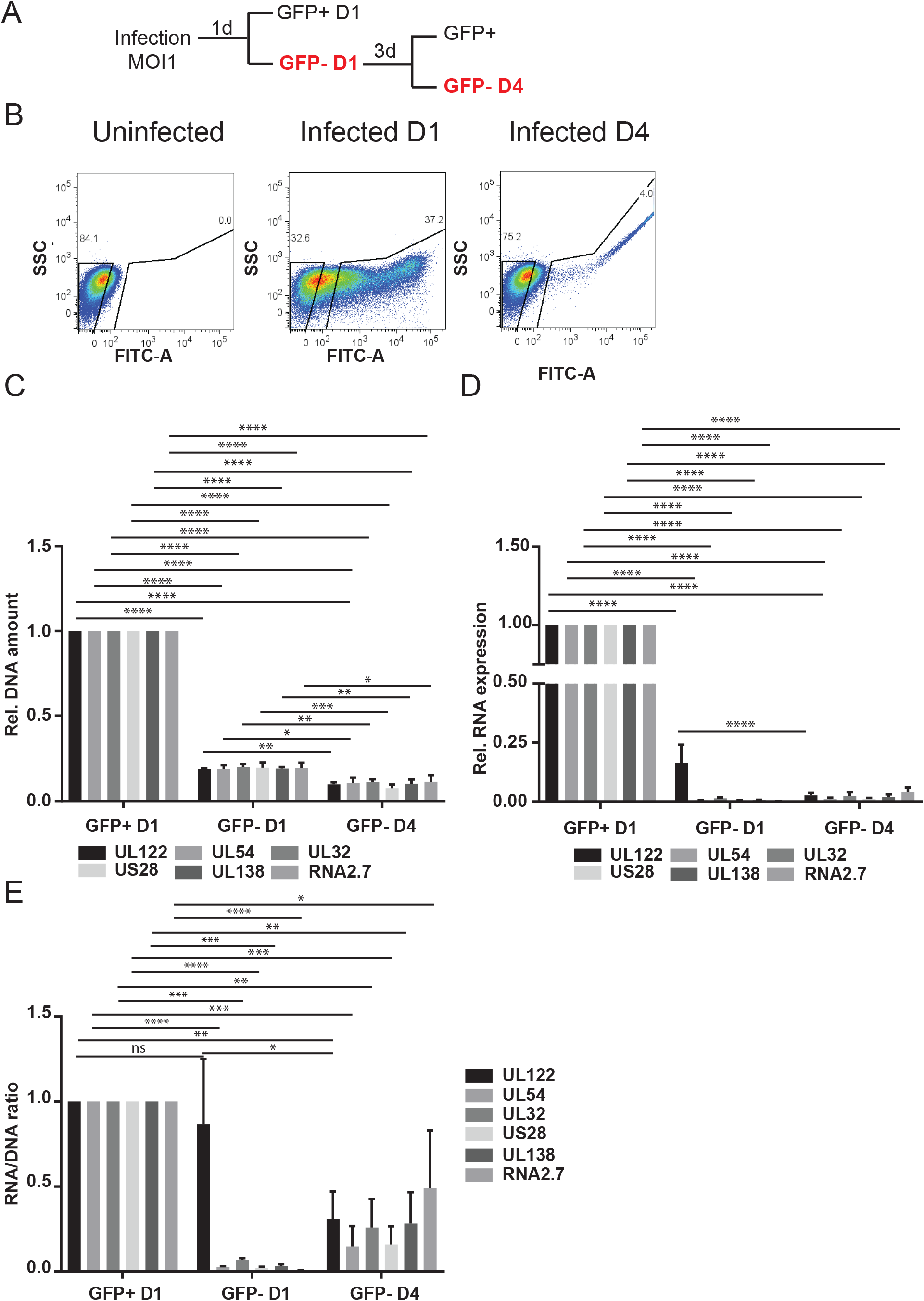
Analysis of GFP-infected cells at 1 and 4 dpi. (A) Schematic outlining the sorting strategy to purify GFP-cells at D1 and D4 post infection. (B) Representative FACS analysis of GFP sorting at D1 and D4 post infection. (C) DNA amount, (D) mRNA expression, (E) RNA/DNA ratio of UL122, UL54, UL32, US28, UL138, and RNA2.7 analyzed at the indicated times post infection and expressed relative to D1. RNA expression values were normalized to GAPDH and the DNA amount was normalized to RNAseP. The statistical significance was calculated by 2-way ANOVA with a Tukey’s multiple comparison test (n=3). The error bars represent the SEM and the asterisks indicate p-values (* P≤ 0.05; ** P≤ 0.01; *** P≤ 0.001; **** P ≤ 0.0001) calculated by comparison of the different genes at each time point.

We found that some GFP-cells become GFP+ by 4 dpi (Fig. 2B). Expression of all classes of viral RNA and DNA copy increased in these cells from 1 to 4 dpi, and then declined after 7 dpi (Fig. S2). In order to determine whether there was a subset of cells in the GFP-population containing quiescent viral DNA, we re-sorted these cells at 4 dpi into GFP-positive and -negative cells for analysis of viral DNA and RNA (Fig. 2A and B). The results show that viral DNA was detectable in the GFP-population at 4 dpi (Fig. 2C, GFP-D4). Analysis of RNA expression showed that some expression of lytic genes was still detectable in the GFP-population at 4 dpi (Fig. 2D,GFP-D4). We conclude that *i)* the GFP reporter is regulated as an early gene in TB40/E*wt*-GFP infected Kasumi-3 cells, as previously noted in fibroblasts (50); *ii)* there is heterogeneity in the kinetics of viral gene expression after infection of Kasumi-3 cells, with activation of early and late gene expression delayed in some cells; *iii)* it is not possible to use sorting for GFP expression to determine whether a small minority of Kasumi-3 cells infected with TB40/E*wt*-GFP contain viral genomes, but remain quiescent.

### US28, UL138, and 2.7 kb RNAs are not differentially expressed in latently infected Kasumi-3 cells

A number of studies have analyzed the viral transcriptome in latency, and some of these have indicated that some lytic genes are also specifically expressed in latency, including US28, UL138, and a non-coding 2.7 kb RNA (RNA2.7) (12, 14, 37, 46, 47, 49, 51-53). We analyzed expression of these genes in GFP+ cells over the course of infection in Kasumi-3 cells. RNA2.7 is expressed at very high levels during lytic infection of fibroblasts (54), and at early times after infection of Kasumi-3 cells (Fig. 3A). UL138 and US28 RNAs are also relatively abundant transcripts during lytic infection. To analyze repression of these genes relative to a lytic gene, we analyzed UL32, a gene that is expressed at levels similar to UL138 and US28 at 4 dpi. Our results show that expression of UL138, US28, and RNA2.7 followed the same pattern as that observed for UL32, with a peak of expression at 4 dpi, followed by a significant decrease at 7 dpi (Fig. 3A). When we analyzed the RNA/DNA ratio, we found that expression of these genes was repressed over the course of infection, with kinetics similar to that of UL32 (Fig. 3C). The RNA/DNA ratio of RNA2.7 was significantly higher than other genes at 4 dpi, consistent with its high level of expression during lytic infection, but there was no difference between UL32 and UL138 or US28 at any time point, and there were no differences in the RNA/DNA ratios between UL32 and any of the other genes analyzed after 4 dpi (Fig. 3C). Thus, there was no preferential expression of UL138, US28, or RNA2.7 over the lytic gene UL32 when latency was established.

**Fig. 3.**
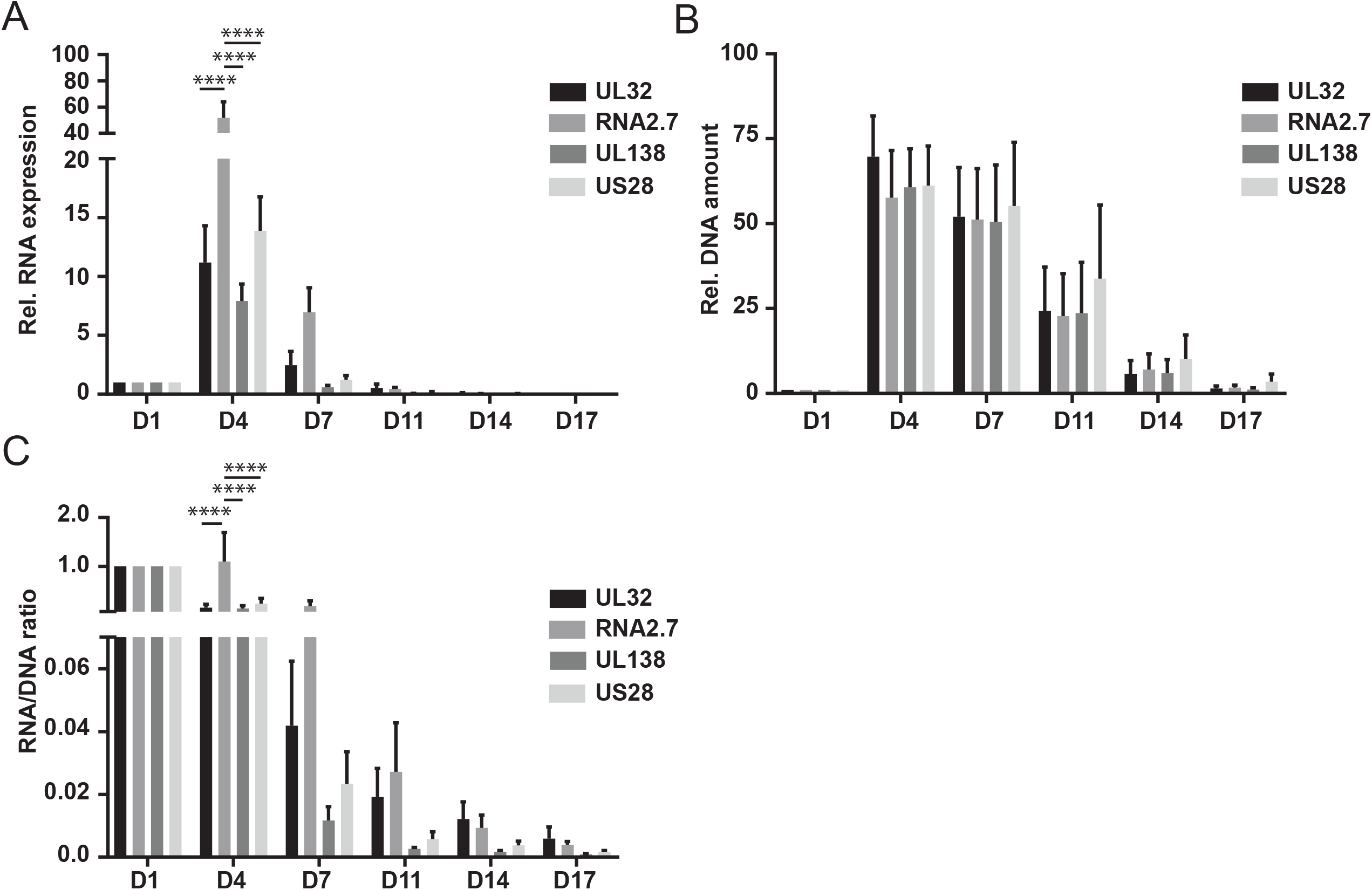
Expression of latency associated RNAs. mRNA expression (A), DNA amount (B) and RNA/DNA ratio (C) of UL32, RNA2.7, UL138 and US28 analyzed at the indicated times post infection and expressed relative to D1. RNA expression values were normalized to GAPDH and the DNA amount was normalized to RNAseP. The statistical significance was calculated by 2way ANOVA with a Tukey’s multiple comparison test (n=3). The error bars represent the SEM and the asterisks indicate p-values (* P≤ 0.05; ** P≤ 0.01; *** P≤ 0.001; **** P ≤ 0.0001) calculated by comparison of the different genes at each time point.

### TNF -α induces reactivation of HCMV in Kasumi-3 cells independently of myeloid cell differentiation

Previous studies showed that TNF-α induced reactivation of HCMV in latently infected Kasumi-3 cells (36). We confirmed this result using slightly different conditions. Kasumi-3 cells were infected with TB40/E*wt*-GFP virus at an MOI of 1-2, flow-sorted for GFP+ cells and cultured for 14 days to establish latency. The cells were then divided into different pools and incubated for a further 3 days in the presence of 5 ng/ml TNF-α or left untreated. Relative to untreated controls, TNF-α induced statistically significant increases in viral DNA copy, RNA expression and the RNA/DNA ratio, as well as production of infectious virus (Fig. 4A and additional RNA analyses of early and late genes in Fig. S3). In addition, we analyzed the percentage of reactivating cells by flow cytometry after staining with an IE1/2-specific antibody. Expression of IE proteins was essentially undetectable in latently infected, untreated Kasumi-3 cells (Fig. 4B, upper right panel). Treatment with TNF-α reproducibly induced expression of IE1/2 proteins in 2-3% of the cells (representative results shown in Fig. 4B, lower left panel).

**Fig. 4.**
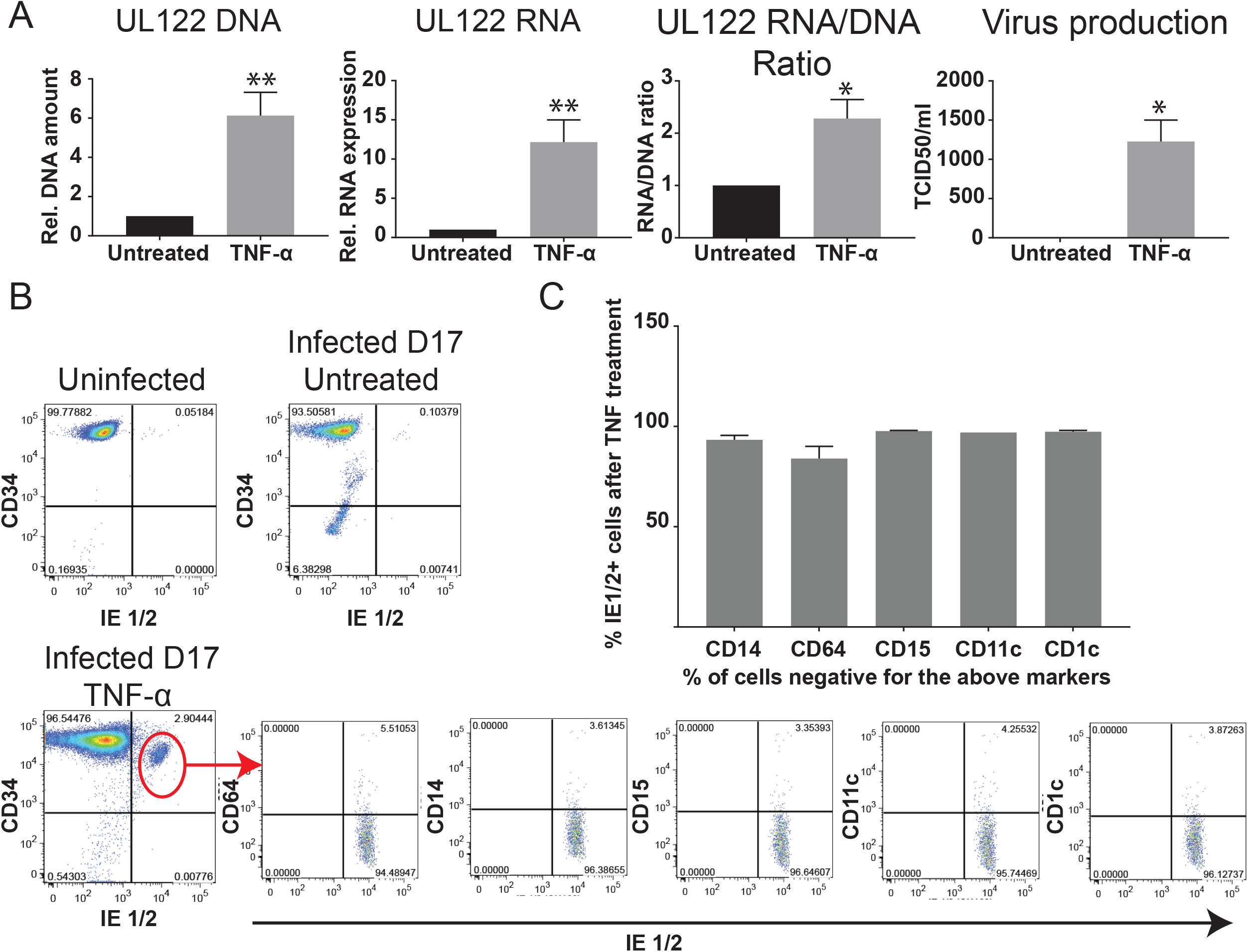
TNF-α induces reactivation independently of differentiation. (A) Latently infected cells (14 dpi) were treated or not with TNF-α for 3 days. On day 17, DNA and RNA from infected cells was analyzed for DNA copy number and expression of viral genes as decribed in Materials and Methods. The release of viral particles in the supernatant was measured by a TCID_50_ assay. The statistical significance was calculated by unpaired *t-test* (n=4). The error bars represent the SEM and the asterisks indicate p-values (* P<0.05; ** P< 0.01). (B) Representative FACS analysis of uninfected, latently infected cells and TNF-α treated cells for the expression of the hematopoietic progenitor marker CD34 and the viral immediate early proteins IE1/2. Reactivating cells were immunophenotyped by gating on the CD34+/IE1/2+ population and analyzing expression of markers of myeloid differentiation (CD64, CD14, CD15, CD11c and CD1c). (C) Percentage of undifferentiated cells within the IE1/2 expressing population in the TNF-α treated cells (CD14-, CD64-, CD15-, CD11c-and CD1c-). The error bars represent the SEM calculated from 3 experiments.

Differentiation of myeloid progenitor cells to a dendritic cell phenotype has long been associated with transcriptional activation of HCMV IE gene expression and reactivation of infectious virus (13, 20, 34, 40). Transformed cell lines are generally resistant to differentiation, and therefore it seemed unlikely that differentiation would play a role in this model. However, the fact that reactivation occurred in only a small percentage of latently infected Kasumi-3 cells left open the possibility that reactivation occurred in a small number of cells with the capacity to differentiate. To investigate this question, we used immunophenotyping to characterize the differentiation state of Kasumi-3 cells carrying reactivating virus. Latently infected TNF-α treated or untreated cells were stained for IE1/2 antibody in combination with the hematopoietic progenitor marker CD34, and for markers of various stages of myeloid differentiation, including CD64, a marker of granulo-monocytic lineage commitment (55), CD14 (monocytes), CD15 (granulocytes), and CD1c and CD11c, which identify major subpopulations of human myeloid dendritic cells (56). Uninfected Kasumi-3 cells are homogeneous with respect to CD34 expression, and do not express other markers of myeloid cell differentiation (Fig. 4B, upper left and Fig. S4). Expression of CD34 was lost in a small percentage of cells after establishment of latency, but latent cells did not differentiate (Fig. S4B). All the latently infected cells were negative for expression of IE proteins (Fig. 4B, upper right panel). 2-3% of the cells became IE+ after treatment with TNF-α, and these cells retained expression of CD34 (Fig. 4B, lower left panel). We then gated on the IE+/CD34+ population to analyze expression of differentiation markers in reactivating cells. ~95% of the IE+/CD34+ cells were negative for CD64- and for markers of mature monocytes and dendritic cells (Fig. 4C). Thus, our data shows that differentiation was not required for transcriptional reactivation of IE gene expression in this model.

### TNF-α induces activation of NF-κB and a DNA damage response in Kasumi-3 cells

Binding of TNF-α to the TNFR1 receptor activates signaling cascades, which lead to activation of transcription factors that control activation of the HCMV Major Immediate Early Promoter (MIEP), including canonical NF-κB (p65/p50) (57-59). Because the IE genes are transcriptionally silent in latently infected cells and they are required for activation of lytic replication, activation of the MIEP is thought to be a requisite first step in reactivation of the virus. Activation of NF-κB is controlled at multiple levels, including cellular localization and post-translational modification at multiple sites (60). TNF/TNFR1-mediated activation of IKK leads to degradation of the inhibitory IκB subunit, which permits translocation of the active p65/p50 complex from the cytoplasm into the nucleus. In addition, IKK mediates phosphorylation of p65 S536 (61, 62), which leads to enhanced transactivation through increased CBP/p300 binding and acetylation of p65 K310 (63). We therefore analyzed total levels of p65, p-p65 S536, and the ratio of p-p65 S536/total p65 in both uninfected and latently infected Kasumi-3 cells treated with or without TNF-α (Fig. 5A). Our results show that all of these increased following treatment with TNF-α in latently infected cells.

**Fig. 5.**
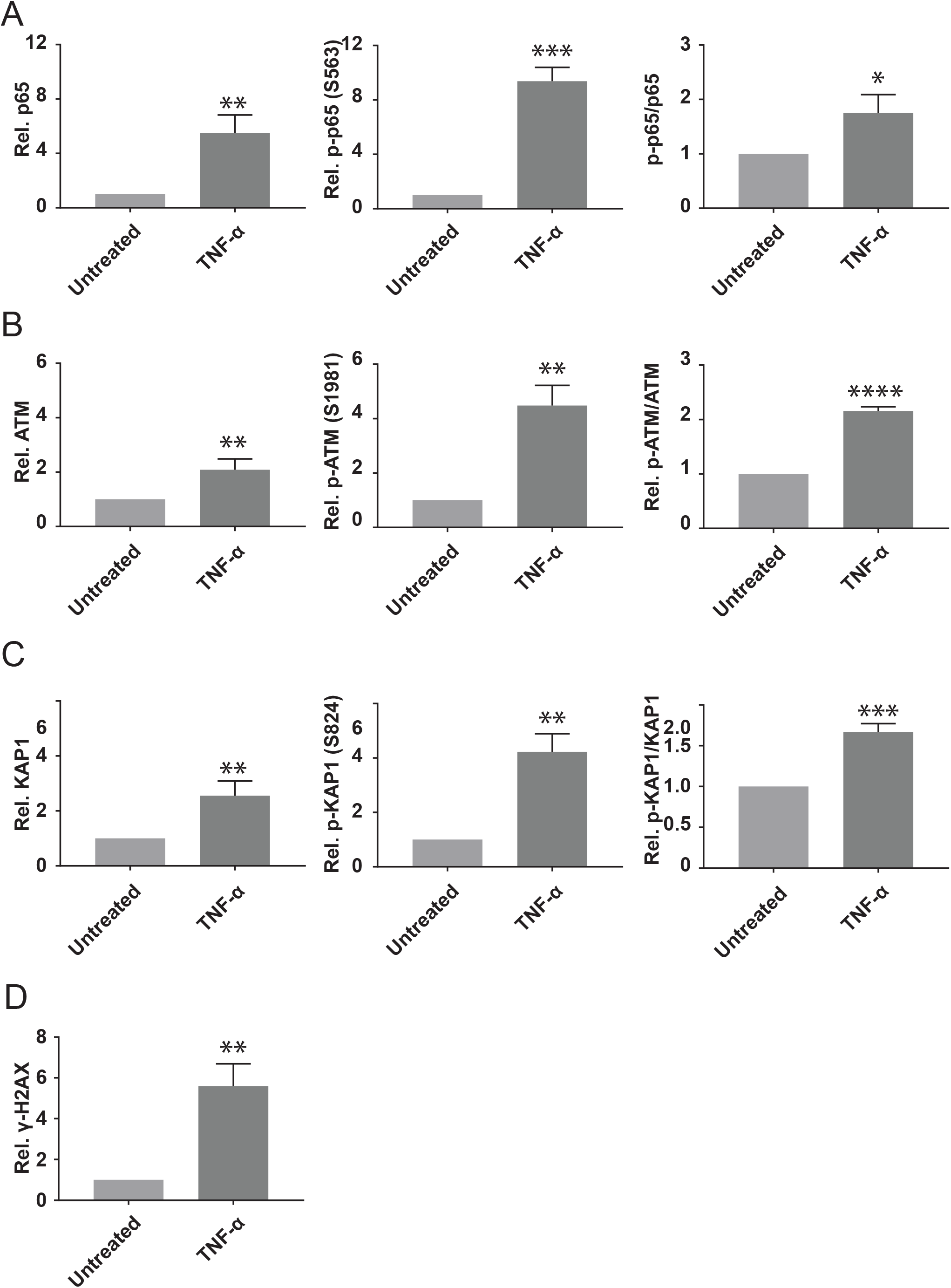
TNF-α mediated reactivation is correlated with activation of NF κ B and ATM signaling. Latently infected cells (14 dpi) were treated for 3 days with or without TNF-*α* 5ng/ml. (A to D) Flow cytometry analysis were performed to assess the activation of NF *κ* B (p65) and DDR (ATM, KAP1 and *γ* H2AX). The activation of p65, ATM and KAP1 and are expressed as the Ratio of Phosphoprotein and total protein compared to untreated cells (p-p65-S536/p65, pATM-S1981/ATM, and p-Kap-S824/Kap1). *γ* H2AX is expressed as ratio of MFI compared to untreated cells. The error bars represent the SEM calculated from 3 independent experiments (* P≤ 0.05; ** P≤ 0.01; *** P≤ 0.001).

Previous studies showed that TNF-α induces activation of ATM (64), and that activation of ATM is sufficient to induce reactivation of HCMV in primary hematopoietic progenitor cells through phosphorylation of KAP-1 (S824) bound to latent viral genomes (40). We therefore analyzed total and phospho-ATM (S1981), total and phospho-KAP-1 (S824) and the ratio of phospho/total proteins. In addition, we analyzed phosphorylation of H2AX (γ-H2AX), a variant histone that is recruited to sites of DNA damage (65). Our results show that all of these markers of DNA damage are increased by treatment of Kasumi-3 cells with TNF-α (Fig. 5B-D).

## DISCUSSION

### Mechanisms of establishment of latency

HCMV establishes latency in myeloid progenitor cells through heterochromatinization and repression of viral gene expression, but the timing and molecular mechanisms that control this process are not well understood. While some studies have shown that viral gene expression is silenced immediately after infection (26, 46), others have shown that viral gene expression is activated in various models at early times post-infection, and that latency is subsequently established in this population of cells (11, 36, 38, 40, 41, 47). Our studies are in agreement with this latter observation. We found that ~90% of infected Kasumi-3 cells become GFP+ at 1 dpi. From 4-8 dpi, expression of all classes of viral genes was activated in this population, viral DNA was amplified, and infectious virus was produced. At later times, viral RNA expression and DNA copy number, and the RNA/DNA ratio decreased. Some lytically infected cells die after infection, and the drop in RNA/DNA may be due in part to loss of these cells. However, we observed a significant decrease in RNA/DNA between live GFP+ cells isolated at 1 dpi, prior to robust viral replication, and 17 dpi, when the cells maintain viral DNA, but expression of viral genes is nearly undetectable. This difference is due at least in part to transcriptional repression of viral genes during the transition from a lytic to latent infection.

Thus, we find that, although activation of the MIEP is lower in myeloid progenitor cells than permissive fibroblasts (26), Kasumi-3 cells support the full cycle of lytic replication. In contrast to fibroblasts, where the infection is propagated throughout the culture to kill all the cells, our studies suggest that some infected Kasumi-3 cells are able to shut down viral replication and survive. Our studies are in agreement with previous studies of Kasumi-3 cells, which showed increased expression of lytic RNAs and amplification of viral DNA relative to early times post-infection, and, in some cases, production of infectious virus, followed by transcriptional repression (36, 37, 66). Although Kasumi-3 cells may be more permissive than primary CD34+ HPCs, previous studies have also shown activation of viral gene expression in HPCs prior to establishment of latency (11, 38, 41, 47).

Recent studies indicate that post-transcriptional establishment of latency may be a more general feature of herpesvirus latency (67-70). Previous studies have shown that activation of the immediate early ICP0 promoter was detectable in some, but not all neurons of mice latently infected with HSV (68). Activation of the late gC promoter was also detectable in the acute stage of infection, but not in latently infected mice. This observation suggests that, although some viral genes may be expressed prior to establishment of latency, there is a point of no return, and full replication of the virus is not compatible with survival. Interestingly, these authors also demonstrated activation of the HCMV MIEP prior to establishment of latency in neurons infected with HSV.

We considered the possibility that there may be two routes to establishment of latency: repression prior to activation of gene expression as well as post-transcriptional repression. Our results show that ~10% of the infected cells are GFP-at 1 dpi. We found that there is heterogeneity in the kinetics of viral gene expression, and that some GFP-cells become GFP+, activate the early and late phases of viral gene expression, and amplify viral DNA with delayed kinetics. Previous studies showed that initiation of HCMV gene expression requires that the cells be in the G0 or G1 phase of the cell cycle (71), and heterogeneity in cell cycle phase may account for some of the observed variation in the kinetics of gene expression. The majority of the cells that were GFP-at 1 dpi remain GFP-at 4 dpi. But, because some GFP-cells expressed lytic viral transcripts, we were unable to determine whether there were other cells that retain viral genomes, but do not express viral RNAs. However, since 90% of the infected cells are GFP+ at 1 dpi, these cells would constitute a very small minority of the infected cells.

Our findings have two important implications. First, the observation that viral gene is expression is first turned on, and then turned off, suggests that there is a mechanism for shutting off viral genomes that are actively engaged in transcription. This may be due to expression of viral genes that facilitate establishment of latency (37, 41, 46-49, 66, 72, 73). However, these genes are also expressed in models of lytic infection, and could not act directly to repress viral gene expression, since they are not localized to the nucleus. An attractive hypothesis that would account for the highly restricted tropism for HCMV latency is that there is a host defense response that shuts off viral transcription specifically in myeloid lineage cells.

Second, shedding of HCMV is often observed in healthy, immunocompetent individuals, suggesting that the virus reactivates with high frequency (74). The finding that HCMV establishes latency in cells following activation of transcription raises the possibility that these genomes may retain a memory of prior activation, and may therefore be poised to reactivate readily under the appropriate conditions. Interestingly, previous studies have shown that HCMV chromatin is specifically marked by H3 lysine 4 methylation following replication of viral DNA in fibroblasts (75). It is tempting to speculate that there may be cell type-specific differences in histone modifications bound to post-replicative viral DNA that affect the potential to establish latency and to reactivate.

### Viral transcription in latency

Previous studies showed that some viral genes that are expressed during lytic infection, particularly UL138 and US28, facilitate establishment of latency in myeloid progenitor cells (37, 41, 46-49, 66, 72, 73). However, analysis of expression of these genes in latently infected cells has been contradictory and confusing. Some studies have shown that some genes are selectively expressed in latency (12, 14, 37, 46, 47, 51-53). In contrast, studies using unbiased transcriptome analyses to detect viral transcripts in both experimental and natural latency, have identified many genes expressed under conditions of latency, and none of these show specific expression of some RNAs in the absence of all other lytic genes (11, 39, 42, 43, 76). This raises the possibility that detection of these transcripts is due to the presence of a small population of lytically infected cells. With the exception of the noncoding miR-UL148D microRNA (37), most previous studies have not included analyses of the RNA/DNA ratio to demonstrate differences in transcriptional activity between lytic genes and latency-associated lytic genes. With the exception of LUNA (14), there has been no analysis of histones bound to latency-associated genes in latently infected cells to support the idea that they are selectively active. We have investigated expression of US28, UL138, as well as the non-coding RNA2.7 in the Kasumi-3 model. We find that the pattern of expression of these genes parallels that of other lytic genes, including UL122, UL123, UL54, and UL32, and analysis of the RNA/DNA ratio shows that transcription of all of these genes is repressed in latency. We suggest that this type of analysis should be included in future studies to demonstrate latency-specific expression of a gene. Collectively, our studies and others are consistent with the idea that these genes facilitate establishment of latency through changes in the relative abundance of viral transcripts, but that they are shut off when latency is established. This situation would be analogous to that of EBV latency, where the EBNA genes are expressed in naive B cells, but are shut off in memory B cells, which are the long term site of latency (77) and HSV, where ICP0 is initially expressed to turn on LAT expression and facilitate heterochromatinization of viral genomes (67).

### What is the role of myeloid cell differentiation in reactivation of HCMV?

Previous studies using primary CD34-derived or monocyte-derived dendritic cells (MoDCs) showed that LPS-induced reactivation of latent HCMV required differentiation to a dendritic cell phenotype (9, 13, 20, 30, 32, 52, 78). Reactivation in immature dendritic cells in response to LPS occurred through induction of IL-6 and activation of ERKs and mitogen and stress-activated kinases (MSK) (52, 79). However, the factors that were used to differentiate dendritic cells in these models are also mediators of inflammation, and it is therefore difficult to distinguish the roles of a general inflammatory response, which could occur in many cell types, from a dendritic cell-specific mechanism of reactivation. In contrast, Kasumi-3 cells, like many transformed cell lines, are refractory to the normal cues that induce differentiation, and are therefore more suitable for studies seeking to dissect the roles of these different processes. Our studies show that differentiation is not required for reactivation in response to TNF-α.

Previous studies showed that, in latently infected primary CD34+ hematopoietic progenitor cells, viral genomes are bound to KAP-1, and that KAP-1 is required to establish latency through heterochromatinization and transcriptional silencing (40). Reactivation could be induced by treatment of the cells with chloroquine, an agent that induces phosphorylation of KAP-1 through activation of ATM, or by knock-down of KAP-1, and this effect was potentiated by co-treatment with TNF-α. These cells retained expression of CD34, suggesting that differentiation was not required for reactivation in this model, although expression of dendritic cell markers was not analyzed (40).

ATM is the master regulator of the response to double-stranded DNA breaks (DSB) (80). ATM is recruited to damage sites by the Mre11-Rad50-Nbs1 (MRN) complex, where it phosphorylates a variety of substrates that facilitate DNA repair, including KAP-1. KAP-1 is a multifunctional protein whose activity is regulated through post-translational modifications (81). The sumoylated form of KAP-1 acts as a repressor of transcription through recruitment of factors that mediate H3K9 de-acetylation/methylation. ATM-mediated phosphorylation of KAP-1 leads to changes in its interaction partners that result in de-condensation of chromatin to allow repair of damaged sites, and this is especially important in repair of breaks that occur in regions of heterochromatin (81-83). The results of Rauwel, et al. (40) therefore suggest that reactivation of HCMV can be induced through chromatin remodeling induced by activation of a DNA damage response independently of differentiation to dendritic cells. Co-treatment with TNF-α potentiated reactivation in response to chloroquine, suggesting that inflammatory mediators that activate NF-κB can act synergistically with chromatin remodeling to induce reactivation of HCMV. Our studies showed that TNF-α was sufficient to induce reactivation in Kasumi-3 cells without inducing differentiation, and that this correlated with activation of NF-κB, as well as ATM and KAP-1. Further studies will be required to fully understand the roles of differentiation, inflammation, and chromatin remodeling in reactivation of HCMV. However, the results of our studies, as well as those of others, suggest that the mechanisms of reactivation of HCMV are not necessarily limited to a dendritic cell specific pathway, but may be induced by more general pathways of inflammation and cellular damage.

## MATERIALS AND METHODS

### Cells and reagents

Kasumi-3 and MRC-5 cells were obtained from the ATCC. Human recombinant TNF-α was obtained from PreproTech (#300-01A). Fluorescent conjugated monoclonal antibodies used in this study are listed in Table 1. TB40/E*wt*-GFP (48) viral stocks were produced and titered as described in supplementary methods.

**TABLE 1.** Antibodies used in this study.

| Marker | Clone | Supplier |
| --- | --- | --- |
| CD34 | 581 | BD Biosciences |
| CD45 | HI30 | BD Biosciences |
| CD1c | F10/21A3 | BD Biosciences |
| CD11c | B-ly6 | BD Biosciences |
| CD14 | MOP9 | BD Biosciences |
| CD15 | HI98 | BD Biosciences |
| CD64 | 10.1 | BD Biosciences |
| IE1/2 | CH160 | Virusys |
| NF-kB p65 | L8F6 | Cell Signaling Technology #9460 |
| phospho-NF-kB p65 (Ser536) | 93H1 | Cell Signaling Technology |
| ATM | D2E2 | Cell Signaling Technology #2873 |
| phospho-ATM (S1981) | D25E5 | Cell Signaling Technology #13050 |
| KAP-1 | EPR5216 | Abcam #109287 |
| phospho-KAP-1 (S824) | EPR5248 | Abcam #133440 |
| γ-H2AX (S139) | EP854(2)Y | Abcam #195189 |

### Infection of Kasumi-3 cells

Kasumi-3 cells were infected with HCMV strain TB40/E*wt*-GFP (48) (MOI 1-2) using centrifugal enhancement of infection, as described in supplementary methods (84). GFP+ and GFP-cells were sorted on a FACSAria-6 laser cell sorter (BD Biosciences). Infected cells were cultured for 13 days to establish latency, with media changes every 2-3 days. For reactivation studies, cells were treated with 5ng/ml TNF-α at 14 dpi for 3 days at 37°C. A TCID_50_ assay was used to determine the infectious titer of infected cell supernatants.

### RNA and DNA extraction

DNA was isolated with the Arcturus pico Pure DNA extraction kit (Thermo Fisher Scientific) as recommended by the manufacturer. RNA was purified with the Direct-zol RNA mini prep kit (Zymo Research) with a DNAse digestion step following manufacturer’s instructions.

### Real time PCR analysis

For all analyses, mock-infected cells were analyzed in parallel with infected cells. These samples were uniformly positive for cellular genes, but negative for viral genes. TaqMan assays specific for viral genes were custom-designed by Life Technologies (Table 2). Relative viral DNA and RNA quantity was analyzed by the 2-(ΔΔCt) method, using pre-designed assays (Life Technologies) for RNaseP (catalog number 4403326) or GAPDH, respectively, as the normalization controls. This method was validated by determining that the efficiencies of the target and reference genes were approximately equal (85) as described in supplementary methods (Fig. S5 and S6). Previous studies have shown that GAPDH is an appropriate normalization control for RNA analysis of CMV-infected cells (86), and we verified that GAPDH Ct values are stable following infection of Kasumi-3 cells (Fig. S7). TaqMan gene expression assays purchased from Life Technologies were used for analysis of cellular RNAs (Hs00174128_m1 for TNF-α and Hs99999905_m1 for GAPDH).

**TABLE 2.**
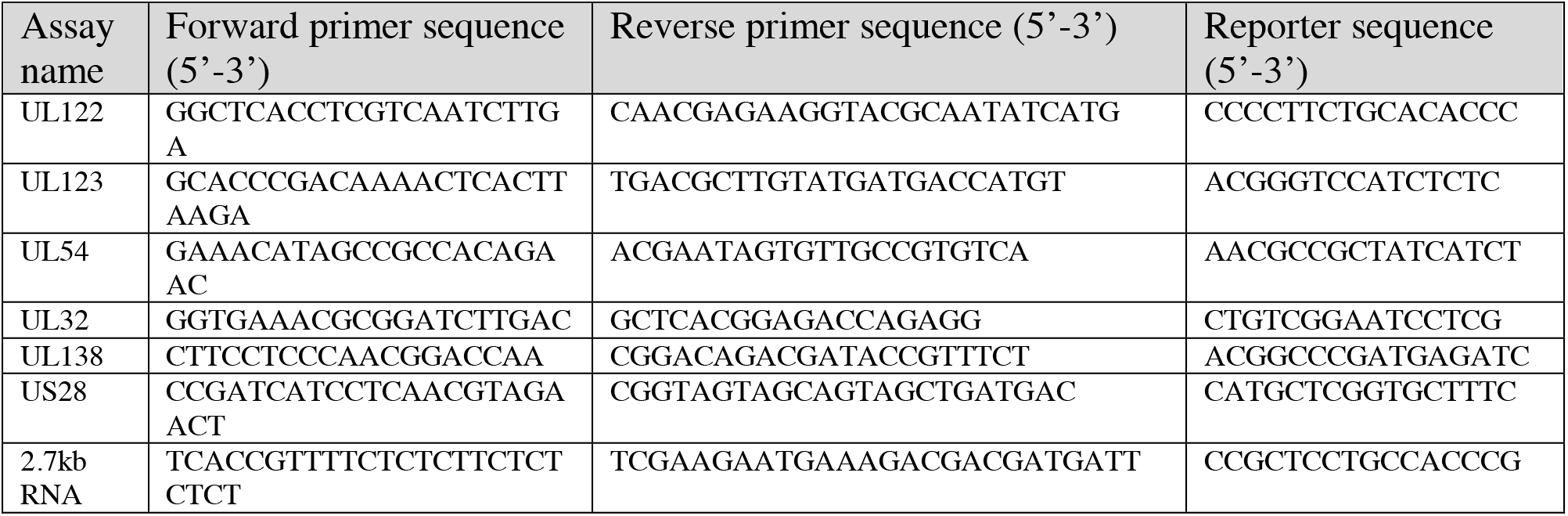
TaqMan assays for analysis of viral DNA and RNA

### Immunophenotyping

Cells were stained with antibodies specific for CD34, CD64, CD14, CD15, CD11c, CD1c and IE 1/2 (CH160) (Table 1) as described in supplementary methods. Data on live cells was acquired on 6-laser BD LSRFortessa SORP flow cytometer and analyzed using FlowJo software (version 9).

### Cytokine stimulation and intracellular phospho-protein analysis

Kasumi-3 cells were treated or untreated with 5 ng/ml TNF for 30’ at 37°C, fixed, permeabilized and washed according to previously described protocols (87, 88). Cells were stained with antibodies specific for p65 (NF-κB), p-p65 (pNF-κB), ATM, p-ATM, KAP-1, p-KAP-1 or γ-H2AX (Table 1). Antibodies were validated using published protocols (89) as described in supplementary methods and Fig. S8. Data were acquired on BD LSRFortessa SORP flow cytometer and analyzed using FlowJo (version 9). The fraction of responding cells for each population was determined as described in supplementary methods as previously described (88, 89).

## ACKNOWLEDGEMENTS

This work was supported by grant P01AI112522 to Michael Abecassis and R01 AI083281 to Scott Terhune from the National Institute of Allergy and Infectious Diseases and by a Cancer Center Support Grant (NCI CA060553) to the Northwestern University Flow Cytometry Core Facility. Cell sorting was performed on a BD FACSAria SORP system, purchased through the support of NIH 1S10OD011996-01. The authors thank Katie Cataldo for preparation of virus stocks, Paul Mehl and Carolina Ostiguin for cell sorting, Dr. Robert Kalejta for the suggestion to analyze RNA/DNA ratios during establishment of latency, and Dr. Michael Abecassis for critical discussions and support. The authors have no conflicts of interest to disclose.

## SUPPLEMENTARY FIGURES

**Fig. S1. Analysis of viral RNA and DNA over the course of infection in Kasumi-3 cells.**

RNAs and DNAs from representative immediate early, early and late genes were analyzed by RT-qPCR and qPCR, respectively, at various times post-infection as described in Fig. 1 and Materials and Methods.

**Fig. S2. Kinetic analysis of HCMV DNA and RNA expression in GFP-cells isolated at 1 dpi.**

(A-C) Relative mRNA expression and DNA amount of UL122, UL54 and UL32 analyzed by RT-qPCR and qPCR, respectively, at the indicated times post infection and expressed relative to D1. Statistical significance was calculated by a one-way ANOVA with a Dunnett’s multiple comparison test (n=3). The error bars represent the SEM and the asterisks indicate p-values (* P≤ 0.05; ** P≤ 0.01; *** P≤ 0.001; **** P ≤ 0.0001*) calculated by comparison to the peak at D4.

**Fig. S3. TNF induces expression of HCMV early and late genes.**

RNAs from the experiments shown in Fig. 6 were analyzed for relative expression of the early gene UL54 and the late gene UL32.

**Fig. S4. Phenotyping of uninfected and latently infected Kasumi-3 cells.**

Representative FACS analysis of uninfected (A) and latently infected (B) cells for the expression of the hematopoietic progenitor marker CD34 and markers of myeloid differentiation (CD64, CD14, CD15, CD11c and CD1c).

**Fig. S5. Analysis of the efficiencies of amplification of viral genes versus RNaseP.**

Viral genes and the cellular gene RNaseP were amplified in samples prepared from serial dilutions of DNA isolated from lytically infected MRC-5 fibroblasts. The ΔCt values (C_T_viral gene – C_T_RNaseP) for each dilution were calculated and plotted against the log ng DNA.

**Fig. S6. Analysis of the efficiencies of amplification of viral RNAs versus GAPDH.**

Viral RNAs and cellular GAPDH RNA were amplified in samples prepared from serial dilutions of cDNA prepared from RNA isolated from lytically infected MRC-5 fibroblasts. The ΔCt values (C_T_viral gene – C_T_GAPDH) for each dilution were calculated and plotted against the log ng cDNA.

**Fig. S7. Validation of GAPDH as a normalization control in HCMV-infected Kasumi-3 cells.**

Data shows average Ct values ± SD for GAPDH at various times after infection. N=4.

**Fig. S8. Antibody staining validation.**

(A) Representative flow cytometric analysis of HeLa cells, untreated (red) or treated with hTNF-α (20ng/ml) and Calyculin A (100nM) for 15 mins (blue), using Phospho-NF-*k*B p65 (Ser563) rabbit mAb and total NF-*k*B p65. (B and C) Representative flow cytometric analysis of HCT116 treated with 200nM NCS, using phospho-ATM (S1981), phospho-KAP1 (S824) mAb, ATM and total KAP1 mAb (blue) compared to untreated control cells (red).

## REFERENCES

1. Eid AJ, Razonable RR. 2010. New developments in the management of cytomegalovirus infection after solid organ transplantation. Drugs 70:965–981.

2. Gilbert GL, Hayes K, Hudson IL, James J. 1989. Prevention of transfusion-acquired cytomegalovirus infection in infants by blood filtration to remove leucocytes. Neonatal Cytomegalovirus Infection Study Group. Lancet 1:1228–1231.

3. Hahn G, Jores R, Mocarski ES. 1998. Cytomegalovirus remains latent in a common precursor of dendritic and myeloid cells. Proc Natl Acad Sci U S A 95:3937–3942.

4. Mendelson M, Monard S, Sissons P, Sinclair J. 1996. Detection of endogenous human cytomegalovirus in CD34+ bone marrow progenitors. J Gen Virol 77 (Pt 12):3099–3102.

5. Sindre H, Tjoonnfjord GE, Rollag H, Ranneberg-Nilsen T, Veiby OP, Beck S, Degre M, Hestdal K. 1996. Human cytomegalovirus suppression of and latency in early hematopoietic progenitor cells. Blood 88:4526–4533.

6. Zhuravskaya T, Maciejewski JP, Netski DM, Bruening E, Mackintosh FR, St Jeor S. 1997. Spread of human cytomegalovirus (HCMV) after infection of human hematopoietic progenitor cells: model of HCMV latency. Blood 90:2482–2491.

7. Taylor-Wiedeman J, Sissons JG, Borysiewicz LK, Sinclair JH. 1991. Monocytes are a major site of persistence of human cytomegalovirus in peripheral blood mononuclear cells. Journal of General Virology 72:2059–2064.

8. Bolovan-Fritts CA, Mocarski ES, Wiedeman JA. 1999. Peripheral blood CD14+ cells from healthy subjects carry a circular conformation of latent cytomegalovirus genome. Blood 93:394–398.

9. Minton EJ, Tysoe C, Sinclair JH, Sissons JG. 1994. Human cytomegalovirus infection of the monocyte/macrophage lineage in bone marrow. Journal of Virology 68:4017–4021.

10. Slobedman B, Mocarski ES. 1999. Quantitative analysis of latent human cytomegalovirus. J Virol 73:4806–4812.

11. Goodrum FD, Jordan CT, High K, Shenk T. 2002. Human cytomegalovirus gene expression during infection of primary hematopoietic progenitor cells: a model for latency. Proc Natl Acad Sci U S A 99:16255–16260.

12. Hargett D, Shenk TE. 2010. Experimental human cytomegalovirus latency in CD14+ monocytes. Proc Natl Acad Sci U S A 107:20039–20044.

13. Reeves MB, Lehner PJ, Sissons JG, Sinclair JH. 2005. An in vitro model for the regulation of human cytomegalovirus latency and reactivation in dendritic cells by chromatin remodelling. J Gen Virol 86:2949–2954.

14. Reeves MB, Sinclair JH. 2010. Analysis of latent viral gene expression in natural and experimental latency models of human cytomegalovirus and its correlation with histone modifications at a latent promoter. J Gen Virol 91:599–604.

15. Hertel L, Lacaille VG, Strobl H, Mellins ED, Mocarski ES. 2003. Susceptibility of immature and mature Langerhans cell-type dendritic cells to infection and immunomodulation by human cytomegalovirus. J Virol 77:7563–7574.

16. Rice GP, Schrier RD, Oldstone MB. 1984. Cytomegalovirus infects human lymphocytes and monocytes: virus expression is restricted to immediate-early gene products. Proc Natl Acad Sci U S A 81:6134–6138.

17. Kalejta RF. 2013. Pre-Immediate Early Tegument Protein Functions, p 141–151. *In* Reddehase MJ (ed), Cytomegaloviruses: From Molecular Pathogenesis to Intervention, vol 1. Caister Academic Press, Norfolk, UK.

18. Saffert RT, Kalejta RF. 2006. Inactivating a cellular intrinsic immune defense mediated by Daxx is the mechanism through which the human cytomegalovirus pp71 protein stimulates viral immediate-early gene expression. J Virol 80:3863–3871.

19. Saffert RT, Kalejta RF. 2007. Human cytomegalovirus gene expression is silenced by Daxx-mediated intrinsic immune defense in model latent infections established in vitro. J Virol 81:9109–9120.

20. Reeves MB, MacAry PA, Lehner PJ, Sissons JG, Sinclair JH. 2005. Latency, chromatin remodeling, and reactivation of human cytomegalovirus in the dendritic cells of healthy carriers. Proceedings of the National Academy of Sciences of the United States of America 102:4140–4145.

21. Cuevas-Bennett C, Shenk T. 2008. Dynamic histone H3 acetylation and methylation at human cytomegalovirus promoters during replication in fibroblasts. J Virol 82:9525–9536.

22. Groves IJ, Reeves MB, Sinclair JH. 2009. Lytic infection of permissive cells with human cytomegalovirus is regulated by an intrinsic ‘pre-immediate-early’ repression of viral gene expression mediated by histone post-translational modification. J Gen Virol 90:2364–2374.

23. Nitzsche A, Paulus C, Nevels M. 2008. Temporal dynamics of cytomegalovirus chromatin assembly in productively infected human cells. J Virol 82:11167–11180.

24. Woodhall DL, Groves IJ, Reeves MB, Wilkinson G, Sinclair JH. 2006. Human Daxx-mediated repression of human cytomegalovirus gene expression correlates with a repressive chromatin structure around the major immediate early promoter. J Biol Chem 281:37652–37660.

25. Zalckvar E, Paulus C, Tillo D, Asbach-Nitzsche A, Lubling Y, Winterling C, Strieder N, Mucke K, Goodrum F, Segal E, Nevels M. 2013. Nucleosome maps of the human cytomegalovirus genome reveal a temporal switch in chromatin organization linked to a major IE protein. Proc Natl Acad Sci U S A 110:13126–13131.

26. Saffert RT, Penkert RR, Kalejta RF. 2010. Cellular and viral control over the initial events of human cytomegalovirus experimental latency in CD34+ cells. J Virol 84:5594–5604.

27. Albright ER, Kalejta RF. 2013. Myeloblastic Cell Lines Mimic Some but Not All Aspects of Human Cytomegalovirus Experimental Latency Defined in Primary CD34+ Cell Populations. J Virol 87:9802–9812.

28. Penkert RR, Kalejta RF. 2010. Nuclear localization of tegument-delivered pp71 in human cytomegalovirus-infected cells is facilitated by one or more factors present in terminally differentiated fibroblasts. J Virol 84:9853–9863.

29. Coronel R, Takayama S, Juwono T, Hertel L. 2015. Dynamics of Human Cytomegalovirus Infection in CD34+ Hematopoietic Cells and Derived Langerhans-Type Dendritic Cells. J Virol 89:5615–5632.

30. Huang MM, Kew VG, Jestice K, Wills MR, Reeves MB. 2012. Efficient human cytomegalovirus reactivation is maturation dependent in the Langerhans dendritic cell lineage and can be studied using a CD14+ experimental latency model. J Virol 86:8507–8515.

31. Riegler S, Hebart H, Einsele H, Brossart P, Jahn G, Sinzger C. 2000. Monocyte-derived dendritic cells are permissive to the complete replicative cycle of human cytomegalovirus. J Gen Virol 81:393–399.

32. Taylor-Wiedeman J, Sissons P, Sinclair J. 1994. Induction of endogenous human cytomegalovirus gene expression after differentiation of monocytes from healthy carriers. J Virol 68:1597–1604.

33. Reeves M, Sinclair J. 2013. Epigenetic Regulation of Human Cytomegalovirus Gene Expression: Impact on Latency and Reactivation, p 330–346. *In* Reddehase MJ (ed), Cytomegaloviruses: From Molecular Pathogenesis to Intervention, vol 1. Caister Academic Press, Norfolk, UK.

34. Sinclair J, Reeves M. 2014. The intimate relationship between human cytomegalovirus and the dendritic cell lineage. Front Microbiol 5:389.

35. Asou H, Suzukawa K, Kita K, Nakase K, Ueda H, Morishita K, Kamada N. 1996. Establishment of an undifferentiated leukemia cell line (Kasumi-3) with t(3;7)(q27;q22) and activation of the EVI1 gene. Jpn J Cancer Res 87:269–274.

36. O’Connor CM, Murphy EA. 2012. A myeloid progenitor cell line capable of supporting human cytomegalovirus latency and reactivation, resulting in infectious progeny. J Virol 86:9854–9865.

37. Pan C, Zhu D, Wang Y, Li L, Li D, Liu F, Zhang CY, Zen K. 2016. Human Cytomegalovirus miR-UL148D Facilitates Latent Viral Infection by Targeting Host Cell Immediate Early Response Gene 5. PLoS Pathog 12:e1006007.

38. Goodrum F, Jordan CT, Terhune SS, High K, Shenk T. 2004. Differential outcomes of human cytomegalovirus infection in primitive hematopoietic cell subpopulations. Blood 104:687–695.

39. Rossetto CC, Tarrant-Elorza M, Pari GS. 2013. Cis and trans acting factors involved in human cytomegalovirus experimental and natural latent infection of CD14 (+) monocytes and CD34 (+) cells. PLoS Pathog 9:e1003366.

40. Rauwel B, Jang SM, Cassano M, Kapopoulou A, Barde I, Trono D. 2015. Release of human cytomegalovirus from latency by a KAP1/TRIM28 phosphorylation switch. Elife 4.

41. Zhu D, Pan C, Sheng J, Liang H, Bian Z, Liu Y, Trang P, Wu J, Liu F, Zhang CY, Zen K. 2018. Human cytomegalovirus reprogrammes haematopoietic progenitor cells into immunosuppressive monocytes to achieve latency. Nat Microbiol 3:503–513.

42. Shnayder M, Nachshon A, Krishna B, Poole E, Boshkov A, Binyamin A, Maza I, Sinclair J, Schwartz M, Stern-Ginossar N. 2018. Defining the Transcriptional Landscape during Cytomegalovirus Latency with Single-Cell RNA Sequencing. MBio 9.

43. Cheung AK, Abendroth A, Cunningham AL, Slobedman B. 2006. Viral gene expression during the establishment of human cytomegalovirus latent infection in myeloid progenitor cells. Blood 108:3691–3699.

44. Banchereau J, Steinman RM. 1998. Dendritic cells and the control of immunity. Nature 392:245–252.

45. Becher B, Tugues S, Greter M. 2016. GM-CSF: From Growth Factor to Central Mediator of Tissue Inflammation. Immunity 45:963–973.

46. Lee SH, Albright ER, Lee JH, Jacobs D, Kalejta RF. 2015. Cellular defense against latent colonization foiled by human cytomegalovirus UL138 protein. Sci Adv 1:e1501164.

47. Goodrum F, Reeves M, Sinclair J, High K, Shenk T. 2007. Human cytomegalovirus sequences expressed in latently infected individuals promote a latent infection in vitro. Blood 110:937–945.

48. Umashankar M, Petrucelli A, Cicchini L, Caposio P, Kreklywich CN, Rak M, Bughio F, Goldman DC, Hamlin KL, Nelson JA, Fleming WH, Streblow DN, Goodrum F. 2011. A novel human cytomegalovirus locus modulates cell type-specific outcomes of infection. PLoS Pathog 7:e1002444.

49. Goodrum F. 2016. Human Cytomegalovirus Latency: Approaching the Gordian Knot. Annu Rev Virol 3:333–357.

50. Qin Q, Penkert RR, Kalejta RF. 2013. Heterologous viral promoters incorporated into the human cytomegalovirus genome are silenced during experimental latency. J Virol 87:9886–9894.

51. Beisser PS, Laurent L, Virelizier JL, Michelson S. 2001. Human cytomegalovirus chemokine receptor gene US28 is transcribed in latently infected THP-1 monocytes. J Virol 75:5949–5957.

52. Reeves MB, Compton T. 2011. Inhibition of inflammatory interleukin-6 activity via extracellular signal-regulated kinase-mitogen-activated protein kinase signaling antagonizes human cytomegalovirus reactivation from dendritic cells. J Virol 85:12750–12758.

53. Keyes LR, Hargett D, Soland M, Bego MG, Rossetto CC, Almeida-Porada G, St Jeor S. 2012. HCMV protein LUNA is required for viral reactivation from latently infected primary CD14(+) cells. PLoS One 7:e52827.

54. Gatherer D, Seirafian S, Cunningham C, Holton M, Dargan DJ, Baluchova K, Hector RD, Galbraith J, Herzyk P, Wilkinson GW, Davison AJ. 2011. High-resolution human cytomegalovirus transcriptome. Proc Natl Acad Sci U S A 108:19755–19760.

55. Olweus J, Lund-Johansen F, Terstappen LW. 1995. CD64/Fc gamma RI is a granulo-monocytic lineage marker on CD34+ hematopoietic progenitor cells. Blood 85:2402–2413.

56. Collin M, McGovern N, Haniffa M. 2013. Human dendritic cell subsets. Immunology 140:22–30.

57. Bhatt D, Ghosh S. 2014. Regulation of the NF-kappaB-Mediated Transcription of Inflammatory Genes. Front Immunol 5:71.

58. Oeckinghaus A, Hayden MS, Ghosh S. 2011. Crosstalk in NF-kappaB signaling pathways. Nat Immunol 12:695–708.

59. Meier JL, Stinski MF. 2013. Major Immediate-Early Enhancer and Its Gene Products, p 152–173. *In* Reddehase MJ (ed), Cytomegaloviruses: From Molecular Pathogenesis to Intervention, vol 1. Caister Academic Press, Norfolk, UK.

60. Chaturvedi MM, Sung B, Yadav VR, Kannappan R, Aggarwal BB. 2011. NF-kappaB addiction and its role in cancer: ‘one size does not fit all’. Oncogene 30:1615–1630.

61. Buss H, Dorrie A, Schmitz ML, Hoffmann E, Resch K, Kracht M. 2004. Constitutive and interleukin-1-inducible phosphorylation of p65 NF-{kappa}B at serine 536 is mediated by multiple protein kinases including I{kappa}B kinase (IKK)-{alpha}, IKK{beta}, IKK{epsilon}, TRAF family member-associated (TANK)-binding kinase 1 (TBK1), and an unknown kinase and couples p65 to TATA-binding protein-associated factor II31-mediated interleukin-8 transcription. J Biol Chem 279:55633–55643.

62. Sakurai H, Chiba H, Miyoshi H, Sugita T, Toriumi W. 1999. IkappaB kinases phosphorylate NF-kappaB p65 subunit on serine 536 in the transactivation domain. J Biol Chem 274:30353–30356.

63. Chen LF, Williams SA, Mu Y, Nakano H, Duerr JM, Buckbinder L, Greene WC. 2005. NF-kappaB RelA phosphorylation regulates RelA acetylation. Mol Cell Biol 25:7966–7975.

64. Choudhary S, Rosenblatt KP, Fang L, Tian B, Wu ZH, Brasier AR. 2011. High throughput short interfering RNA (siRNA) screening of the human kinome identifies novel kinases controlling the canonical nuclear factor-kappaB (NF-kappaB) activation pathway. J Biol Chem 286:37187–37195.

65. Lukas J, Lukas C, Bartek J. 2011. More than just a focus: The chromatin response to DNA damage and its role in genome integrity maintenance. Nat Cell Biol 13:1161–1169.

66. Humby MS, O’Connor CM. 2015. Human Cytomegalovirus US28 Is Important for Latent Infection of Hematopoietic Progenitor Cells. J Virol 90:2959–2970.

67. Raja P, Lee JS, Pan D, Pesola JM, Coen DM, Knipe DM. 2016. A Herpesviral Lytic Protein Regulates the Structure of Latent Viral Chromatin. MBio 7.

68. Proenca JT, Coleman HM, Connor V, Winton DJ, Efstathiou S. 2008. A historical analysis of herpes simplex virus promoter activation in vivo reveals distinct populations of latently infected neurones. J Gen Virol 89:2965–2974.

69. Proenca JT, Coleman HM, Nicoll MP, Connor V, Preston CM, Arthur J, Efstathiou S. 2011. An investigation of herpes simplex virus promoter activity compatible with latency establishment reveals VP16-independent activation of immediate-early promoters in sensory neurones. J Gen Virol 92:2575–2585.

70. Proenca JT, Nelson D, Nicoll MP, Connor V, Efstathiou S. 2016. Analyses of herpes simplex virus type 1 latency and reactivation at the single cell level using fluorescent reporter mice. J Gen Virol 97:767–777.

71. Spector DH. 2013. Exploitation of Host Cell Cycle Regulatory Pathways by HCMV, p 247–263. *In* Reddehase MJ (ed), Cytomegaloviruses: From Molecular Pathogenesis to Intervention, vol 1. Caister Academic Press, Norfolk, UK.

72. Lee SH, Caviness K, Albright ER, Lee JH, Gelbmann CB, Rak M, Goodrum F, Kalejta RF. 2016. Long and Short Isoforms of the Human Cytomegalovirus UL138 Protein Silence IE Transcription and Promote Latency. J Virol 90:9483–9494.

73. Petrucelli A, Rak M, Grainger L, Goodrum F. 2009. Characterization of a novel Golgi apparatus-localized latency determinant encoded by human cytomegalovirus. J Virol 83:5615–5629.

74. Wylie KM, Mihindukulasuriya KA, Zhou Y, Sodergren E, Storch GA, Weinstock GM. 2014. Metagenomic analysis of double-stranded DNA viruses in healthy adults. BMC Biol 12:71.

75. Nitzsche A, Steinhausser C, Mucke K, Paulus C, Nevels M. 2012. Histone H3 lysine 4 methylation marks postreplicative human cytomegalovirus chromatin. J Virol 86:9817–9827.

76. Cheng S, Caviness K, Buehler J, Smithey M, Nikolich-Zugich J, Goodrum F. 2017. Transcriptome-wide characterization of human cytomegalovirus in natural infection and experimental latency. Proc Natl Acad Sci U S A 114:E10586–E10595.

77. Thorley-Lawson DA, Hawkins JB, Tracy SI, Shapiro M. 2013. The pathogenesis of Epstein-Barr virus persistent infection. Curr Opin Virol 3:227–232.

78. Kew VG, Wills MR, Reeves MB. 2017. LPS promotes a monocyte phenotype permissive for human cytomegalovirus immediate-early gene expression upon infection but not reactivation from latency. Sci Rep 7:810.

79. Kew VG, Yuan J, Meier J, Reeves MB. 2014. Mitogen and stress activated kinases act co-operatively with CREB during the induction of human cytomegalovirus immediate-early gene expression from latency. PLoS Pathog 10:e1004195.

80. Stracker TH, Roig I, Knobel PA, Marjanovic M. 2013. The ATM signaling network in development and disease. Front Genet 4: 37.

81. Iyengar S, Farnham PJ. 2011. KAP1 protein: an enigmatic master regulator of the genome. J Biol Chem 286:26267–26276.

82. Ziv Y, Bielopolski D, Galanty Y, Lukas C, Taya Y, Schultz DC, Lukas J, Bekker-Jensen S, Bartek J, Shiloh Y. 2006. Chromatin relaxation in response to DNA double-strand breaks is modulated by a novel ATM- and KAP-1 dependent pathway. Nat Cell Biol 8:870–876.

83. Goodarzi AA, Noon AT, Deckbar D, Ziv Y, Shiloh Y, Lobrich M, Jeggo PA. 2008. ATM signaling facilitates repair of DNA double-strand breaks associated with heterochromatin. Mol Cell 31:167–177.

84. Umashankar M, Goodrum F. 2014. Hematopoietic long-term culture (hLTC) for human cytomegalovirus latency and reactivation. Methods Mol Biol 1119:99–112.

85. Livak KJ, Schmittgen TD. 2001. Analysis of relative gene expression data using realtime quantitative PCR and the 2(-Delta Delta C(T)) Method. Methods (Duluth) 25:402–408.

86. Watson S, Mercier S, Bye C, Wilkinson J, Cunningham AL, Harman AN. 2007. Determination of suitable housekeeping genes for normalisation of quantitative real time PCR analysis of cells infected with human immunodeficiency virus and herpes viruses. Virol J 4:130.

87. Chow S, Minden MD, Hedley DW. 2006. Constitutive phosphorylation of the S6 ribosomal protein via mTOR and ERK signaling in the peripheral blasts of acute leukemia patients. Exp Hematol 34:1183–1191.

88. Marvin J, Swaminathan S, Kraker G, Chadburn A, Jacobberger J, Goolsby C. 2011. Normal bone marrow signal-transduction profiles: a requisite for enhanced detection of signaling dysregulations in AML. Blood 117:e120–130.

89. Krutzik PO, Nolan GP. 2003. Intracellular phospho-protein staining techniques for flow cytometry: monitoring single cell signaling events. Cytometry A 55:61–70.

